# Genome expansion and lineage-specific genetic innovations in the world’s largest organisms (*Armillaria*)

**DOI:** 10.1101/166231

**Authors:** György Sipos, Arun N. Prasanna, Mathias C. Walther, Eoin O’Connor, Balázs Bálint, Krisztina Krizsán, Brigitta Kiss, Jaqueline Hess, Torda Varga, Jason Slot, Robert Riley, Bettina Bóka, Daniel Rigling, Kerrie Barry, Juna Lee, Sirma Mihaltcheva, Kurt Labutti, Anna Lipzen, Rose Waldron, Nicola M. Moloney, Christoph Sperisen, László Kredics, Csaba Vágvölgyi, Andrea Patrigniani, David Fitzpatrick, István Nagy, Sean Doyle, James B. Anderson, Igor V. Grigoriev, Ulrich Güldener, Martin Münsterkötter, László G Nagy

## Abstract

*Armillaria* species are both devastating forest pathogens and some of the largest terrestrial organisms on Earth. They forage for hosts and achieve immense colony sizes using rhizomorphs, root-like multicellular structures of clonal dispersal. Here, we sequenced and analyzed genomes of four *Armillaria* species and performed RNA-Seq and quantitative proteomic analysis on seven invasive and reproductive developmental stages of A. ostoyae. Comparison with 22 related fungi revealed a significant genome expansion in *Armillaria*, affecting several pathogenicity-related genes, lignocellulose degrading enzymes and lineage-specific genes likely involved in rhizomorph development. Rhizomorphs express an evolutionarily young transcriptome that shares features with the transcriptomes of fruiting bodies and vegetative mycelia. Several genes show concomitant upregulation in rhizomorphs and fruiting bodies and shared cis-regulatory signatures in their promoters, providing genetic and regulatory insights into complex multicellularity in fungi. Our results suggest that the evolution of the unique dispersal and pathogenicity mechanisms of *Armillaria* might have drawn upon ancestral genetic toolkits for wood-decay, morphogenesis and complex multicellularity.

---

The genus *Armillaria* includes some of the most devastating forest pathogens worldwide. *Armillaria* causes root rot disease in both gymno-and angiosperms, in forests, parks, and even vineyards in more than 500 host plant species^1^. Most *Armillaria* species are facultative necrotrophs, which, after colonizing and killing the root cambium, transition to a saprobic phase, decomposing dead woody tissues of the host. As saprotrophs, *Armillaria* spp. are white rot (WR) fungi, which can efficiently decompose all components of plant cell walls, including lignin, (hemi-)cellulose and pectin^2^. They produce fleshy fruiting bodies (honey mushrooms) that appear in large clumps around infected plants and produce sexual spores. The vegetative phase of *Armillaria* is predominantly diploid rather than dikaryotic like most basidiomycetes.

Individuals of *Armillaria* can reach immense sizes and include the ‘humongous fungus’, one of the largest terrestrial organisms on Earth^3^, measuring up to 965 hectares and 600 tons^4^ and displaying a mutation rate ⍰ 3 orders of magnitude lower than most filamentous fungi^5^. Individuals reach this immense size via growing rhizomorphs, 1-4 mm wide dark mycelial strings that allow the fungus to bridge gaps between food sources or host plants^1,6^ (hence the name shoestring root rot). Rhizomorphs develop through the aggregation and coordinated parallel growth of hyphae, similarly to some fruiting body tissues^7,8^. As migratory and exploratory organs, rhizomorphs can grow approximately one m/year and cross several meters underground in search for new hosts, although roles in uptake and long-range translocation of nutrients have also been proposed^1,9,10^. Root contact by rhizomorphs is the main mode of infection by the fungus, which makes the prevention of recurrent infection in *Armillaria*-contaminated areas particularly difficult^1^. Despite their huge impact on forestry, horticulture and agriculture, the genetics of the pathogenicity of *Armillaria* species is poorly understood. The only -omics data to date have highlighted a substantial repertoire of plant cell wall degrading enzymes (PCWDE) and secreted proteins, among others, in *A. mellea* and *A. solidipes^11,12^*, while analyses of the genomes of other pathogenic basidiomycetes (e.g. *Moniliophthora^13,14^, Heterobasidion^15^, Rhizoctonia^16^*) identified genes coding for PCWDEs, secreted and effector proteins or secondary metabolism (SM) as putative pathogenicity factors. However, the life cycle and unique dispersal strategy of *Armillaria* prefigure other evolutionary routes to pathogenicity, which, along with potential other genomic factors (e.g. transposable elements^17^) are not yet known.

Here, we investigate genome evolution and the origins of pathogenicity in *Armillaria* using comparative genomics, transcriptomics and proteomics. We sequenced the genomes of four *Armillaria* species to combine with that of related saprotrophic, hemibiotrophic and mycorrhizal fungi. Transcript and proteome profiling of invasive and reproductive developmental stages shed light on the role of rhizomorphs, several putative pathogenicity factors as well as the morphogenetic mechanisms of rhizomorph and fruiting body development.

## Results

We report the genomes of *A. ostoyae, A. cepistipes, A. gallica* and *A. solidipes* sequenced using a combination of PacBio and Illumina technologies. Genomes of *Armillaria* species were assembled to 103–319 scaffolds comprising 58–85 Mb and were predicted to contain 20,811– 25,704 genes (Table 1). In comparison, other sequenced species of the Physalacriaceae, *Flammulina velutipes* and *Cylindrobasidium torrendii* have 12,218 (35.6Mb) and 13,940 (31.5Mb) genes, respectively, while the sister genus of *Armillaria*, *Guyanagaster necrorhiza* has 14,276 (53.6Mb)(Fig. 1). *Armillaria* species share significant synteny, comprising macro-to microsynteny (Supplementary Figure 1), whereas mesosynteny, characteristic of certain fungal groups^18^ was not observed. Transposable element content of *Armillaria* genomes shows a modest expansion relative to other Agaricales and even distribution along the scaffolds, suggesting that their genome expansion is not driven by transposon proliferation, as observed in other plant pathogens^17^ (Supplementary Figs. 2-3, Supplementary Table 1).

**Table 1.** Summary statistics of the new genomes reported in this paper.

| Species | Assembly size<br>(# scaffolds) | N50 | # genes | TE content | BUSCO | CEGMA |
| --- | --- | --- | --- | --- | --- | --- |
| *A. cepistipes* | 75.5 (182) | 8 (3.29 Mb) | 23461 | 34.8% | 95.1% | 97.9% |
| *A. gallica* | 85.34 (319) | 22 (1.04 Mb) | 25704 | 32.6% | 98.6% | 99.8% |
| *A. ostoyae* | 60.1 (106) | 9 (2.28 Mb) | 22705 | 27.3% | 95.7% | 99.2% |
| *A. solidipes* | 58.01 (229) | 21 (716 kb) | 20811 | 20.9% | 98.4% | 99.8% |

**Fig. 1.**
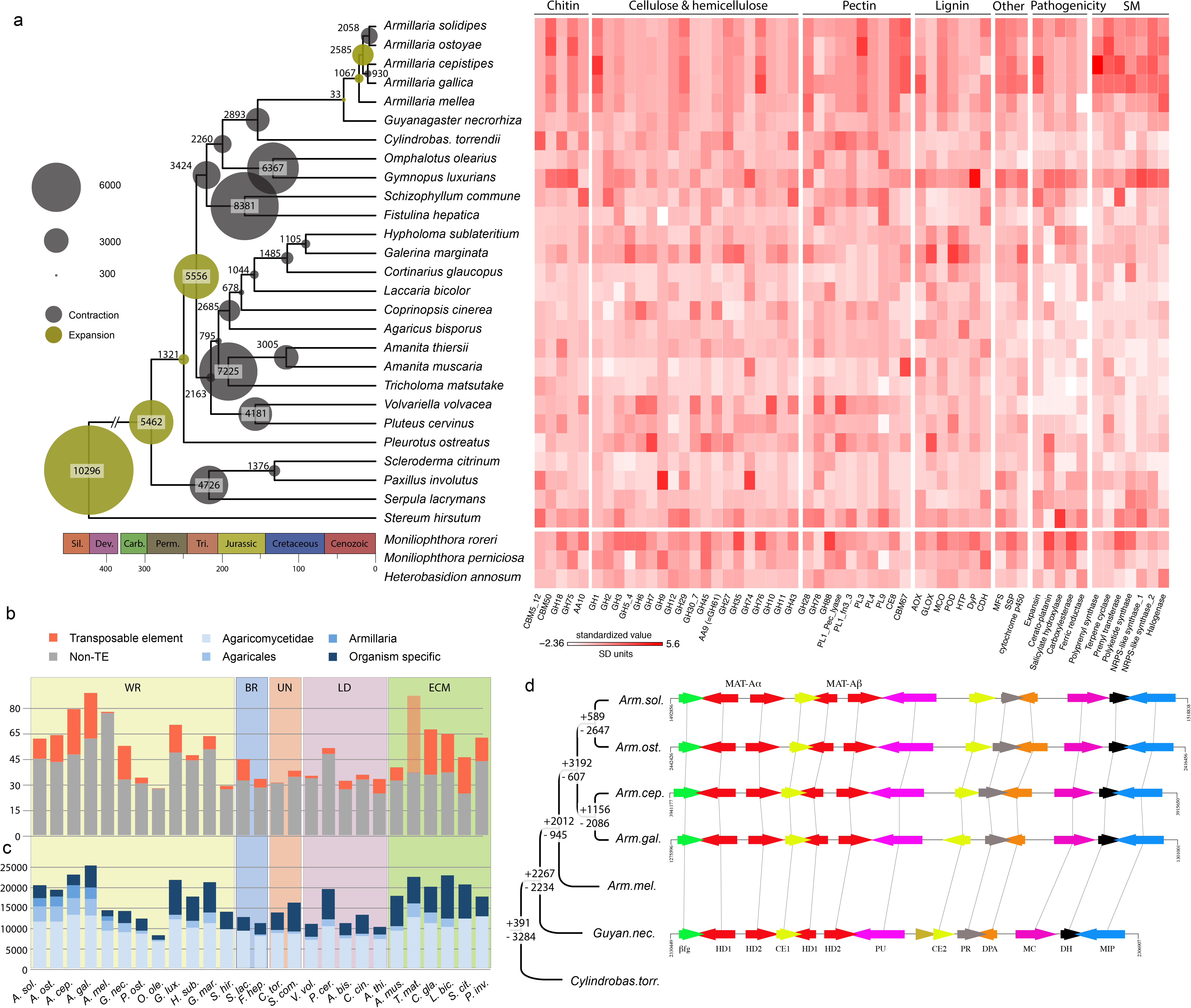
Genome evolution and phylogenomics in *Armillaria*. (a) Reconstructed gene duplication/loss histories along a time-calibrated phylogeny of 27 fungal species shows the expansion of protein coding gene repertoires in *Armillaria* spp. The heatmap shows gene copy numbers for PCWDE and pathogenicity-related gene families in the 27 species examined. (b) Assembly sizes (in Mb) for 27 species broken down by TEs (orange) and non-TEs (gray). Data for *A. mellea* are incomplete due to the highly fragmented available assembly (29,300 scaffolds). *Tricholoma matsutake* has been truncated for clarity. See Supplementary Fig. 2 for details (c) Numbers of predicted genes by phylogenetic conservation. (d) Numbers of duplicated (+) and lost (-) genes inferred for the Physalacriaceae, showing the net genome expansion in *Armillaria* spp. and the structure of their mating loci showing high levels of synteny. The locus comprises of four homeodomain (HD) transcription factors arranged into *MAT-A*α and *MAT-A*β respectively. *β-fg* and *MIP* genes anchor the locus. Genes containing carbohydrate esterase (CE), proline racemase (PR), deoxyribose-phosphate aldolase (DPA), metacapases (MC) and dehydrogenase/reductase (DH) domains (pfam) are located within the locus. For clarity, *A. mellea* is not displayed as it’s assembly is highly contiguous.

Phylogenomic analysis based on 835 conserved single copy genes (188,895 amino acid sites) confirmed the position of *Armillaria* in the Physalacriaceae, with *Guyanagaster* and *Cylindrobasidium* as their closest relatives (Fig. 1/a). We estimated the age of pathogenic *Armillaria* spp. at 21 million years (myr) and their divergence from *Guyanagaster* at 42 myr (Supplementary Fig. 4), coincident with decreasing temperatures and the spread of deciduous forests in the Eocene. Reconstruction of genome-wide gene duplication and loss histories in 27 Agaricales species revealed an early origin for most genes, followed by lineage-specific gene losses in most family and genus level groups, except *Armillaria*, which showed a net genome expansion: 16,687 protein-coding genes were inferred for the most recent common ancestor (mrca) of *Armillaria* (2,012 duplications, 945 losses), as opposed to 14,720 and 14,687 for the mrca of *Armillaria* and *Guyanagaster* and that of *Armillaria*, *Guyanagaster* and *Cylindrobasidium*, respectively (Fig. 1/a,d, Supplementary Fig. 5). Further expansion to 19,272 genes was inferred for the mrca of *A. solidipes, A. ostoyae, A. cepistipes* and *A. gallica* (3,192 duplications, 607 losses), although the highly fragmented *A. mellea* assembly might cause some duplications to map to this instead of the preceding node.

Duplicated genes were enriched in functions related to chitin and cellulose binding, polysaccharide metabolism (peroxidase, lyase, hydrolase and oxidoreductase activity), peptidase activity, transmembrane transport, extracellular region and gene expression regulation (p<0.05, hypergeometric test, HT, Supplementary Table 2). In line with this, 214 domains were significantly overrepresented in *Armillaria* genomes (p<0.05, HT) relative to other agarics, including several peptidase, glycoside hydrolase (GH) and pectinase domains, CBM50-s, expansins, multicopper oxidases as well as several pfams related to secondary metabolism (Supplementary Table 3). We found 571 protein clusters specific to *Armillaria* and *Guyanagaster* or subclades therein (Supplementary Table 4). These included CE4 chitooligosaccharide deacetylases, CBM50-s (LysM), iron permeases (FTR1), and 19 transcription factor (TF) families, among others, although a significant proportion (70%) had no functional annotations. Taken together, these results suggest that gene family expansion was the predominant mode of genome evolution in *Armillaria* and that the observed genome expansion is largely concerned with diverse extracellular functions, including several lineage-specific innovations some of which have previously been associated with pathogenicity.

## Putative pathogenicity-related genes in *Armillaria*

Plant pathogenic fungi possess diverse gene repertoires to invade host plants and modulate their immune system^18–20^. We cataloged 20 families of putative pathogenesis-related genes to assess if *Armillaria* share expansions of these families with other plant pathogens (Supplementary Table 5). *Armillaria* species are enriched in expansins (p=4x10^-5^, Fisher exact test, FET) and possess many cerato-platanin genes, which contribute to unlocking cell-wall polysaccharide complexes and cause cell death in host plants, respectively and might act as a first-line cell-lysis weaponry during invasion (Fig. 1/a). Carboxylesterases and salicylate hydroxylases show moderate enrichment in both *Armillaria* and *Moniliophthora* species, along with other weakly pathogenic taxa (e.g. *Marasmius fiardii*). Salicylate hydroxylases have been implicated in developing a tolerance to salicylic acid, which is released to block the jasmonic acid defense pathway of plants upon infection. In contrast, cutinases (CE5) are missing from *Armillaria* species but present in *Moniliophthora* spp.^14^ that primarily infect through leaves. LysM domains (CBM50) are overrepresented in *Armillaria* compared to other Agaricales (p=2x10^-8^, FET), while GH75 chitosanases are exclusively found in *Armillaria* and *Moniliophthora* species (Fig 1/a, Supplementary Table 5). In other plant pathogens, these are involved in modifying fungal cell wall composition and/or capturing chitin residues to mask chitin-triggered immune signals in host plants^21–25^, suggesting similar roles in *Armillaria* too.

Homologs of SM genes reported from *Heterobasidion^15^* are overrepresented in *Armillaria* species (p=0.03 – 10^-11^, FET, Fig. 1/a, Supplementary Table 3,5). These include terpene cyclases, NRPS-like prenyl transferases, and halogenases, as well as trichodiene and polyprenyl synthases, which have been linked to fungal pathogenesis, virulence and competition with other microbes^26^.

## *Armillaria* species as wood-rotting fungi

It has been shown that plant-fungal interactions in both pathogens^14,15,18^ and mycorrhizal fungi^27–29^ can draw upon the PCWDE repertoire of saprotrophic ancestors. Therefore, we compared the PCWDE complement of *Armillaria* species to that of other Agaricales with diverse lifestyles. In the saprotrophic phase of their lifecycle, *Armillaria* species cause white rot on wood, which is reflected in their PCWDE repertoire. Their genomes encode lignin-, cellulose-, hemicellulose-and pectin-degrading enzymes, indicating the potential to degrade all plant cell wall (PCW) components (Fig. 1/a, Supplementary Table 5). Lignolytic families are underrepresented in *Armillaria* (p=0.02 - 10^-10^, FET), whereas cellulose-and xylan-degrading families generally show similar gene counts to other WR species, with notably higher copy numbers of GH1 (β-glucosidase, p=3x10^-6^, FET). On the other hand, several pectinolytic families are overrepresented in Armillaria. Pectin degrading families include GH28, GH78 and GH88, polysaccharide lyase (PL) 1, 3, 4 and 9 as well as carbohydrate esterase 8 (CE8), of which PL3, CE8 and CBM67 rhamnose binding modules are significantly enriched in *Armillaria* spp. (p=0.02 - 10^-11^, FET) compared to WR Agaricales. The pectinolytic repertoire of *Armillaria* is unusual for WR fungi^30^ and might indicate links to dicot pathogenicity^20^. The PCWDE repertoire of *Armillaria* species underpins their ability to act powerful WR decayers and provides a pool to draw upon as necrotrophic pathogens. It might enable them to gain access to wood and avoid competition with other microbes by damaging live trees, a strategy that is unavailable to most WR fungi.

## Expression profiles of rhizomorphs

*Armillaria* species spread either clonally by rhizomorphs or with sexual spores produced on fruiting bodies. Rhizomorphs are unique multicellular structures of *Armillaria* species, but their functions and morphogenetic origins are debated^10^. We obtained gene expression and proteome profiles for actively growing rhizomorph tips (0-5 cm from the apex) of *A. ostoyae C18* buried in sawdust-rich medium and four fruiting body stages (Supplementary Table 6). Differential expression analysis identified 1,303 and 1,610 over-and underexpressed genes in rhizomorphs relative to vegetative mycelium grown on the same medium (FC_VM_>4, p≤0.0-.05respectively, marking one of the largest expression change in our experiments (Fig. 2). Similarly, the highest number of unique proteins (n=729) was detected in rhizomorphs compared to vegetative mycelia (Fig 2/c-d). Upregulated genes were enriched for GO terms related to carbohydrate, lipid and secondary metabolism, hydrophobins and pectinolysis (Supplementary Table 6). Global expression profiles suggest that rhizomorphs are transitional between vegetative mycelium and fruiting bodies (Supplementary Figs. 6-7). Expression profiles of several PCWDE families and putative pathogenicity-related genes (cerato-platanins, expansins) in rhizomorphs resembles vegetative mycelia (Supplementary Figs 8-12), whereas that of many putative morphogenesis-related genes was shared with fruiting bodies.

**Fig. 2.**
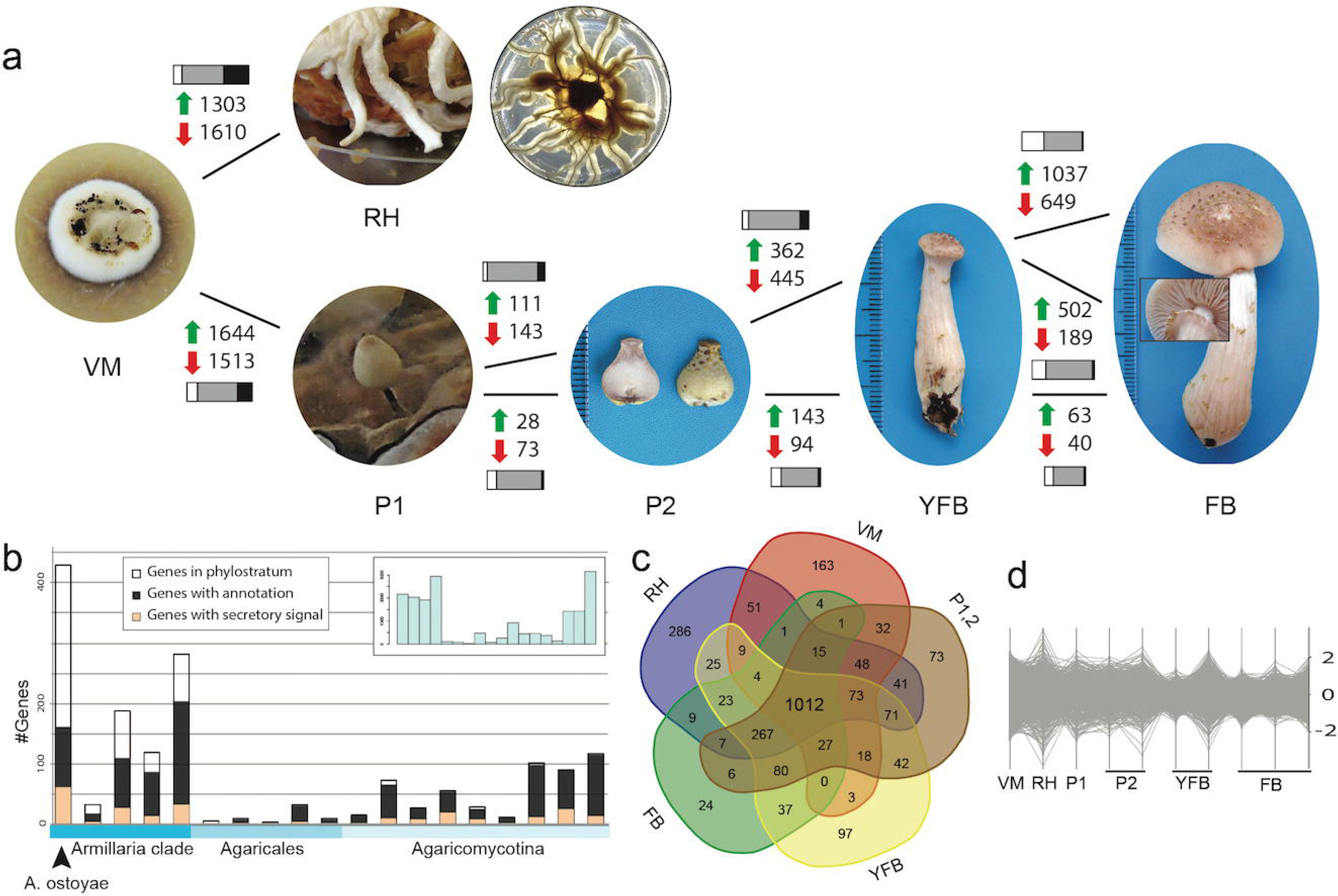
Expression analysis of invasive and reproductive developmental stages of *A. ostoyae C18*. (a) Gene expression and proteomic map of ten developmental stages and tissue types. Green and red arrows denote the numbers of significantly up-and downregulated genes (p<0.05, FC>4), respectively. Bar diagrams show the number of detected proteins common (grey area) to successive developmental stages, and the ones unique to the first (white) or the next (black) stage. The development or rhizomorphs (RH, examples shown for growth in the dark and light, left to right) and that of stage I primordia (P1) from vegetative mycelium (VM) denote the largest expression change, whereas stage II primordia (P2), young fruiting bodies (YFB) and fruiting bodies (FB) show smaller numbers of DEGs (FDR-corrected P<0.05, FC>4). Each sample was sequenced in 3 biological replicates (b) Phylostratigraphic analysis of 1,303 gene upregulated in rhizomorphs. Inset graph shows the age distribution of all genes in the genome of *A. ostoyae*. (c) Number of detected proteins overlapping in the compared developmental stages. Note the similarity between vegetative mycelium and rhizomorphs and between all complex multicellular stages (267 proteins). (d) Protein abundance profiles in ten developmental stages and tissue types. The y-axis shows fold change values.

Rhizomorphs express an evolutionarily young transcriptome, with most upregulated genes specific to *A. ostoyae* or the *Armillaria* clade (including *Guyanagaster* Fig. 2/b). Most of these young genes lacked functional annotations but were conserved in *Armillaria* species, suggesting important functions in the genus. Genes belonging to older phylostrata had comparatively more functional annotation terms. We found 414 genes, which had expression maxima in rhizomorphs and were significantly upregulated relative to vegetative mycelium (FC_VM_≥4, Supplementary Table 7). The most highly upregulated genes included expansins ≥ (log2FC_VM_=10.44), bZip and C_2_H_2_ transcription factors, three caspase genes, hydrophobins, cytochrome p450s, GHs as well as several unannotated genes. We found an overexpression of H_2_O_2_ neutralizing 5-oxoprolinases and signs of intensified biogenesis and cargo of extracellular proteins along the secretory pathway (Supplementary Figs 13-14).

We observed significant overexpression of three cerato-platanins, three expansins, two carboxylesterases, as well as SM-related trichodiene (five genes), polyketide (six genes) synthases and a polyprenyl synthase, among others (Supplementary Table 8). Notably, all expansins showed upregulation in fruiting bodies too, which could indicate a role in fungal cell wall remodeling instead of cellulose degradation, to which fungal expansin-related genes have mostly been linked^31^. A gene coding for a family 6 bacterial extracellular solute-binding protein showed an expression peak in rhizomorphs but low or negligible expression in other developmental stages. Bacterial homologs of this gene function as Fe^3+^ and thiamin transporters and are required for the pathogen’s survival in the host. This gene is a member of an *Armillaria*-specific cluster with homologs in all *Armillaria* species and bacterial proteins as closest BLAST hits (although ML gene trees don’t support HGT). Its Fe^3+^ transporting ability and nearly exclusive expression in rhizomorphs makes us speculate of a role in substrate and/or host exploitation.

A diverse suite of PCWDE encoding genes was expressed at high levels in rhizomorphs, although overall, PCWDE gene expression was higher in vegetative mycelium (Supplementary Figs. 8-11). Pectin-degrading CAZy families showed the most remarkable upregulation, including several GH28s, pectinesterases, a GH88 and PL3 gene (Supplementary Fig. 11), although the latter showed elevated expression throughout fruiting body development also. Similarly, two HTP genes and eight laccases were significantly overexpressed (Supplementary Table 8), consistent with records of laccase activity in rhizomorphs^32^. However, both HTPs and three laccases were also upregulated in fruiting bodies, with HTPs mostly in gill tissues, suggesting a potential role in morphogenesis instead of lignin degradation. Cellulose, hemicellulose and xylan-degrading genes showed highest expression in vegetative mycelium, with the exception of three GH3, a GH43, GH76, AA9 and lignolytic POD genes (Supplementary Fig. 11). Overall, the significant expression of PCWDE genes in rhizomorphs (Supplementary Figs. 8-11) suggest that rhizomorphs may not only explore the surrounding for new substrates, but could actively assimilate nutrients from available food sources. Taken together, the expression data show that under our experimental settings rhizomorphs expressed only a moderate set of pathogenicity-related and PCWDE genes, but which combined could confer invasive properties to rhizomorphs.

## Morphogenesis

The morphogenetic machinery underlying rhizomorph development is among the least known aspects of the biology of *Armillaria*. As multicellular structures, rhizomorphs express a variety of genes encoding cell-wall proteins, including hydrophobins, pore-forming toxins, two CBM67 and four ricin-B-lectins, an annexin and a cell-wall integrity sensor, among others, indicating several fruiting body-like functions, such as hyphal adhesion, communication or defense. We found cell-wall biosynthesis genes to be generally but moderately upregulated in rhizomorphs (and stipes) (Supplementary Table 9, Supplementary Fig. 15). Further, several GMC oxidoreductases, two mating-type pheromone receptors, CBM50-s, a CBM5_12 and chitooligosaccharide deacetylase genes were significantly overexpressed (the latter two also in stipes). Four hydrophobins reached their highest expression in rhizomorphs whereas two showed high expression in rhizomorphs and stipes, but not caps or vegetative mycelium (Supplementary Table 9, Supplementary Fig. 16). A homolog of the *Cryptococcus* red and far-red sensing Tco3 photoreceptor was expressed at high levels (log_2_FC=4.36). Although its function in *Cryptococcus* is unknown^33^, its overexpression in rhizomorphs and in brown film forming mycelia of *Lentinula^34^* could suggest morphogenesis-related functions.

We detected 19 significantly upregulated TFs, ten of which had peak expression in rhizomorphs across our experimental conditions. Although based on global TF expression, rhizomorphs are most similar to vegetative mycelium (Supplementary Fig. 17), several TFs showed shared overexpression in rhizomorphs and fruiting bodies. For example, the expression of AROS_01275 peaked in rhizomorphs and stipes, whereas a zf-Mynd TF was highly upregulated in rhizomorphs and all fruiting body stages, and is thus a candidate for governing complex multicellular development in *A. ostoyae*.

## Fruiting bodies

Fruiting body production is probably the largest morphogenetic transition in the life cycle of Basidiomycota. It involves the reorganization of hyphal growth patterns and the execution of a complex developmental program. Indeed, we detected the largest number of differentially expressed genes (DEGs) and the second largest change in proteome in Stage I primordia, including 1644 up-and 1513 downregulated genes (FC_VM_>4, p>=0.05, Supplementary Table 10) and 435 unique proteins. Upregulated genes were enriched for gene expression regulation, lipid metabolism, amino acid transport and ribonuclease activity (Fig. 2/a, Supplementary Table 9) and included ten TFs and two Dicer-like genes. Following stage I primordia, we tracked expression levels in two cell lineages. While the stipe lineage showed minor changes throughout development (<150 DEGs, <300 unique proteins), cap differentiation included up to 1037 DEGs and 646 unique proteins related to signal transduction, carbohydrate metabolism or the regulation of biological processes (Supplementary Table 10). Genes upregulated in gills (n=502, 56 unique proteins) were enriched for functions related to protein phosphorylation, carbohydrate metabolism, protein kinase activity and ribonucleotide binding.

Hydrophobin genes showed stage and tissue-specific expression patterns, with four genes significantly overexpressed in stipes, one in stage I primordia and stage II caps and another from stage I primordia through young fruiting bodies, with lower expression values in later stages. Hydrophobins were generally not expressed in cap tissues (Supplementary Fig. 16), suggesting alternative sources of hydrophobin-related functionalities in *Armillaria* caps. An analysis of cysteine-rich, hydrophobic proteins (excluding hydrophobins) revealed 33 and 13 developmentally regulated genes with expression maxima in fruiting bodies and caps, respectively (Supplementary Table 11). In addition, three cell wall galactomannoproteins show high expression in fruiting bodies (up to logFC_VM_=13.5, Supplementary Table 12), two of which were expressed only in caps, whereas AROS_19505 was highly expressed throughout development. These genes are homologous to the *Aspergillus* hydrophobic cell-surface protein HsbA, which has been hypothesized to recruit hydrolytic enzymes during lignocellulose degradation^35^ and appressorium attachment to host surfaces^36^. As lignocellulose-related processes are inactive in fruiting bodies, their expression suggests specific roles during cap development, possibly in cell-wall remodeling. Several chitin metabolism-related genes were upregulated in fruiting bodies, including GH88 chitinases, a CBM50 containing gene, a GH75 and six chitooligosaccharide deacetylases (Supplementary Table 10), which might be related to generating development-specific cell-wall architectures. Interestingly, certain HTPs, GMC oxidoreductases (AA3) and pectinolytic genes, generally linked to lignocellulose degradation, were also upregulated in fruiting bodies (Supplementary Fig. 18).

## Defense-related genes in rhizomorphs and fruiting bodies

Defense against predators and competitors is a fundamental property for complex multicellular structures. We detected a significant proportion of putative defense-related genes of *A. ostoyae* upregulated in rhizomorphs, fruiting bodies or both (Supplementary Table 13). Fruiting bodies expressed three fungal hemolysins: a deuterolysin and a bacterial MTX2 pore-forming toxin gene were highly upregulated across fruiting body development, whereas a second MTX2 homolog showed upregulation in stipes and rhizomorphs. The insecticidal activities of these hemolysins^37^ and the observed expression patterns suggest a role in defense of fruiting body structures. Two cerato-platanin genes were significantly upregulated in rhizomorphs and fruiting bodies, as well as four ricin-B-lectin genes, previously implicated in defense against nematodes and insects^37^. A homolog of the nematocidal *Coprinopsis* ccl2^38^ and a gene encoding a thaumatin-like protein were highly expressed from stage I primordia to young fruiting bodies mature fruiting bodies and vegetative mycelium. Taken together, the upregulated genes indicate a phylogenetically and functionally diverse defense arsenal expressed in both fruiting bodies and rhizomorphs.

## Shared morphogenetic machineries between rhizomorphs and fruiting bodies

Rhizomorphs share a complex multicellular organization with fruiting bodies. Consistent with this, we observed several genes with similar expression patterns in rhizomorphs and various fruiting body tissues (Fig. 3). These included two mating type pheromone receptor and the white collar 1-2 genes, which mediate the initiation of fruiting body development and could point to shared developmental origins of rhizomorphs and fruiting bodies or two CBM67 genes that were upregulated in rhizomorphs and all fruiting body samples relative to vegetative mycelium. There were 442 genes that showed >fourfold elevated expression over vegetative mycelium and relatively constant expression across rhizomorph and fruiting body samples (Fig. 3/a, Supplementary Table 14), suggesting they are linked to complex multicellularity in *A. ostoyae*. A systematic analysis identified 2,225 genes showing higher expression in rhizomorphs and stipes than in vegetative mycelium and corresponding cap tissues, of which 63 were at least fourfold more abundant in all four pairs of successive developmental stages (Fig. 3/b). These included most of the yeast cell wall biosynthesis pathway homologs (Supplementary Table 9, Supplementary Fig. 19), pore forming toxins, hydrophobins, TFs and SM genes, among others. For caps and rhizomorphs 1,728 and 28 such genes were found, including several GMC oxidoreductases, a GH88, a ricin-b-lectin and other SM genes (Fig. 3/c, Supplementary Table 15). This indicates that rhizomorph development extensively draws upon fruiting body genes, and makes us speculate that fruiting body development could have been the cradle for the evolution of rhizomorphs in *Armillaria* spp.

**Fig. 3.**
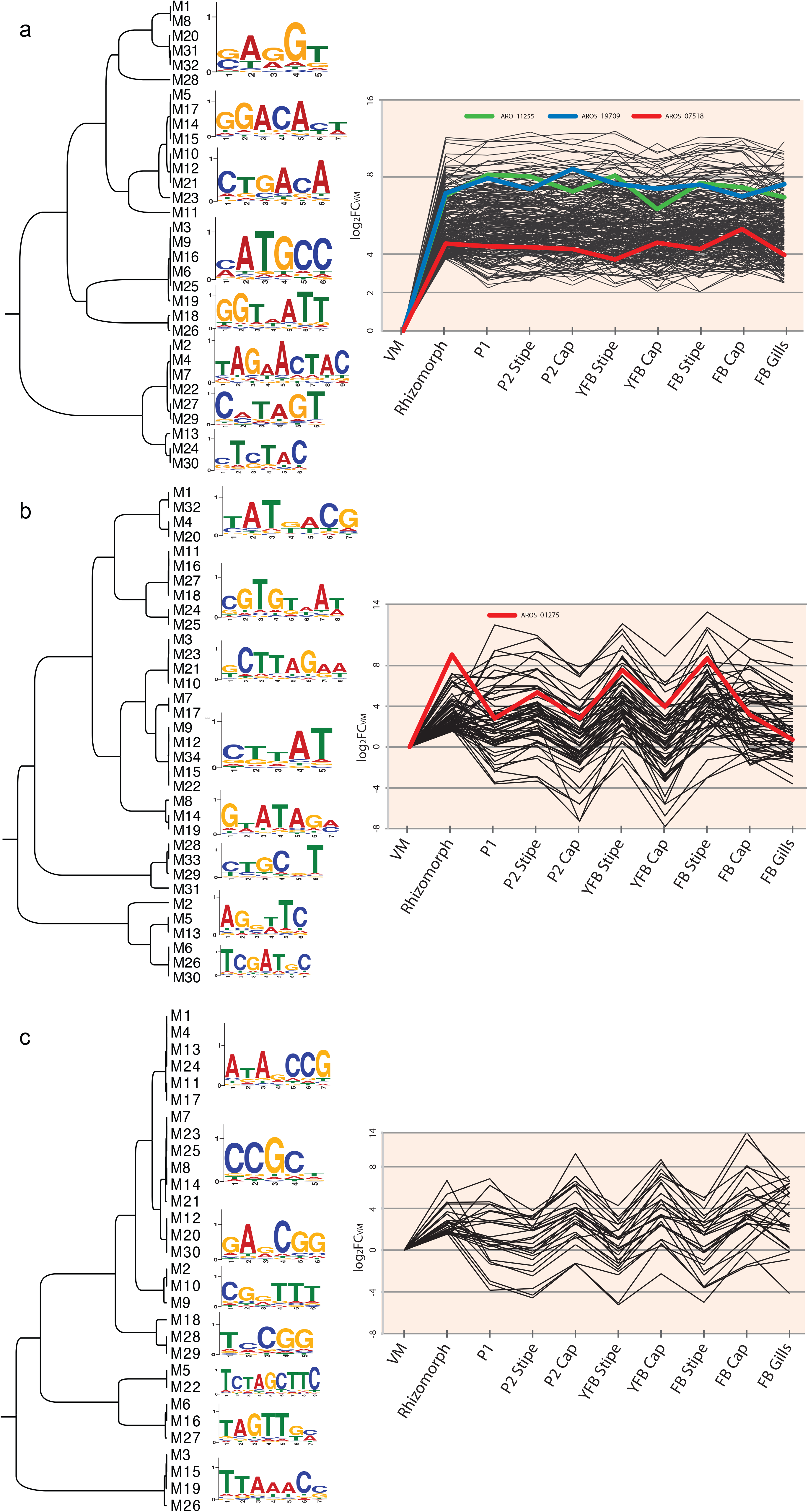
Predicted motifs among promoter regions of co-expressed genes in Armillaria rhizomorphs and fruiting bodies. Putative transcription factor binding sites and expression profiles are shown for three co-expressed gene sets identified in rhizomorph and fruiting body transcriptomes. (a) 442 genes that are at least fourfold upregulated relative to vegetative mycelium and show relatively constant expression in all complex multicellular structures (stage I primordia through fruiting bodies). (b) and (c) 63 and 28 genes with concomitant upregulation in rhizomorphs and stipe tissues and rhizomorphs and caps, respectively, showing over fourfold expression difference in rhizomorphs relative to mycelium and all four pairs of successive developmental stages. Highlighted are putative TF genes detected in the co-expressed gene sets.

To assess whether co-regulated genes possess a common cis-regulatory signature, we searched for putative TF binding sites (TFBS) by *de novo* motif discovery^39^ in 1kb upstream of 63 stipe/rhizomorph and 28 cap/rhizomorph co-expressed genes as well as in promoters of those that showed constant high expression in all complex multicellular structures (Fig. 3). High-scoring motifs in each gene set grouped into eight clades in a UPGMA analysis. To validate these motif predictions, we checked whether the predicted motifs match known consensus TFBS sequences of *Saccharomyces* in the JASPAR^40^ database. All predicted motifs matched experimentally determined yeast TFBS sequences in JASPAR, which, although the phylogenetic distance is large between *S. cerevisiae* and *A. ostoyae*, suggests strong biological relevance to our predictions. Taken together, the detected co-expressed genes and the presence of shared motifs in their promoters could suggest common regulatory mechanisms for rhizomorphs and fruiting bodies and probably represent elements of the regulatory networks governing multicellular development as well as cap and stipe differentiation in *Armillaria*.

## Discussion

Forest pathogens in the genus *Armillaria* evolved from saprotrophic ancestors in the Agaricales. They have unusually large genomes for WR saprotrophs, which evolved mostly by gene family diversification, in contrast with genome expansions in other plant pathogenic fungi, which are primarily driven by TE proliferation^17^. We found many lineage-specific genes (including putative pathogenicity factors and pectinolytic families), coincident overrepresentation of several pathogenicity-related genes with other Basidiomycota (above all *Moniliophthora* species), suggesting convergent origins of pathogenicity in the Agaricales. *Armillaria* species encode a full complement of PCWDE genes, which comes as no surprise given their ability to cause WR, but provides a resource to draw upon in the evolution of pathogenicity. This could give *Armillaria* spp. early access to dead wood and a strategy to bypass competition with other microbes.

Rhizomorphs are some of the most unique structures of *Armillaria* spp. that enable them to become the largest terrestrial organism on Earth^3,5^. They express a wide array of genes involved in secondary metabolism, defense, PCW degradation and to a lesser extent pathogenesis, which indicate active nutrient uptake and adaptations to a soil-borne lifestyle where competition with other microbes and defense against predators are crucial. This underpins their role as exploratory and assimilating organs adapted to bridge gaps between food sources or potential host plants. In terms of morphogenesis, rhizomorphs resemble fruiting bodies, with many stipe-and to a smaller extent cap-upregulated genes showing concomitant upregulation and shared cis-regulatory signatures. Both rhizomorphs and fruiting bodies show key traits of complex multicellularity, such as 3-dimensional organization, cell adhesion or a highly integrated developmental program. We hypothesize that the observed similarity in gene expression indicates common developmental origins and suggest that the evolution of rhizomorphs may have extensively drawn upon the genetic toolkit of fruiting body development in the Agaricomycotina. We identified genes putatively involved in cap-and stipe morphogenesis, as well as some co-expressed in all complex multicellular stages and candidate TF binding motifs within their promoters. These represent putative building blocks of the gene regulatory circuits governing mushroom development and enabled us to zero in on the genetic bases of complex multicellularity in fungi. This study has provided comparative genomics insights into the evolution, pathogenicity and multicellular development of a group of devastating forest pathogens. It should facilitate further understanding of the biology of *Armillaria*, which, combined with new genomic resources and *in planta* interrogation of their pathogenic behavior could bring the development of efficient strategies for containing the spread and damage of *Armillaria* root-rot disease in various forest stands soon within reach.

## Acknowledgements

This work was supported by the ‘Momentum Program’ of the Hungarian Academy of Sciences under LP2014/12, and an OTKA-ERC_HU #118722 from the NRDI Office to L.G.N. Fungal research at Maynooth University is part-funded by a Science Foundation Ireland Investigator award to SD (12/IP/1695). LC-MS facilities were funded by a competitive award from Science Foundation Ireland (12/RI/2346 (3)). G.S. acknowledges support from a WSL grant for a guest researcher in Switzerland and funding from GINOP-2.3.2-15-2016-00052. The work conducted by the U.S. Department of Energy Joint Genome Institute (JGI), a DOE Office of Science User Facility, was supported by the Office of Science of the U.S. Department of Energy under Contract No. DE-AC02-05CH11231. IN was supported by a Janos Bolyai Research Fellowship of the Hungarian Academy of Sciences. We also thank Renate Heinzelmann for verifying the haploid status of the *A. ostoyae* and *A. cepistipes* isolates used for genome sequencing and Catherine Aime and Francis Martin for providing access to unpublished genome data produced by JGI. Two anonymous reviewers are thanked for their constructive comments on earlier versions of the paper.

## Author contributions

G.S. and L.G.N. conceived the study, G.S., K.B., C.S., J.L., U.G., S.M., K.L., A.L., R.W., M.C.W., A.P., I.V.G. and M.M. performed genome sequencing, assembly and annotation, N.M.M., E.O. and S.D. performed the proteomic study, D.F. analyzed mating genes, J.H. performed analyses of TE content, A.P., B.K., I.N., Be.B., L.K., C.V. and B.B. obtained and analyzed RNA-Seq data for *A. ostoyae*, A.P., K.K., J.S., R.R. and L.G.N. performed comparative genomic and phylogenomic analyses, T.V. performed the molecular clock dating. L.G.N, A.P., J.A., G.S. wrote the manuscript. D.R. contributed tested strains for sequencing. All authors read and commented on the manuscript.

## Competing Financial Interests

The authors declare no competing financial interests.

## Methods

### Strains and fungal material used for genome sequencing

The diploid *A. ostoyae* C18 is a field isolate from Switzerland^41^ and the haploid C18/9 is derived from C18 as a single spore isolate. The *A. cepistipes* B5 haploid^42^ was originally isolated from a fruiting body on *Fagus sylvatica* in Italy (H. Marxmüller) and then included in the set of haploid tester strains by J.J. Guillaumin. The *A. ostoyae* C18/9 and *A. cepistipes* B5 haploid isolates were cultured on Roth and Shaw plates (for 1 L RS: 40 g malt extract, 20g dextrose, 5g bacto peptone, 19g agar) covered with cellophane sheets, and incubated at 25°C for three weeks. Prior to genomic DNA extraction, fungal mycelia were detached and harvested from the cellophane sheets, and frozen in liquid nitrogen. The *A. gallica* 21-2 and *A. solidipes* 28-4 strains were maintained on 2% malt extract, 2% agar medium. To produce mycelium for DNA extraction, strains were grown in liquid CYM medium (d-glucose 20g, yeast extract 2 g, peptone 2 g, MgSO_4_.7H_2_O 0.5 g, KH_2_PO_4_ 0.46 g, K_2_HPO_4_ 1g, 1l water) in Petri dishes with 8-10 small pieces of inoculum floated on the air-water interface. Genomic DNA was extracted using the PowerMax MOBIO DNA isolation kit according to the manufacturer’s instructions. Note that Burdsall and Volk (2008) have proposed that the *A. solidipes* and *A. ostoyae* are one species and that the earlier name *A. solidipes* from a North American collection should always be used in place of *A. ostoyae*. For consistency with recent historical usage, however, we choose to consider *A. ostoyae* in Europe as a separate, but closely related species, to *A. solidipes* in North America

We developed an *in vitro* fruiting system for *A. ostoyae* C18 based on the protocol published for *A. mellea^43^*. A modified RST medium, termed RSTO was prepared in 720 ml jars and inoculated with *A. ostoyae* C18, incubated for 28 days at 24°C in the dark, then placed in a growth chamber at 15°C with a 10/14 hour light/dark cycle (light intensity: 11 µE m^-2^s^-1^). Vegetative mycelia, rhizomorphs, stage I and II primordia, young and mature fruiting bodies were harvested as shown on Fig. 1. Fruiting body stages were defined to conform to the general notation^44^ of mushroom developmental stages as closely as possible. Stage I primordia were defined as up to 1-2 mm tall fruiting body initials without a clear differentiation of a pileal section. Stage II primordia had a developed button-shaped cap, up to 7-10 mm tall. Young fruiting bodies were 30-50 mm tall with a developed hymenial cavity but prior to cap expansion. From stage II primordia, caps and stipes were separated and RNA was extracted separately. Mature fruiting bodies were separated to stipe, gills and cap. RNA was extracted by using the RNEasy Midi kit (QIAGEN) following the manufacturer’s instructions. RNA quality was checked by gel electrophoresis and the Agilent 2200 TapeStation before library preparation. Biological triplicates were analyzed for all sample types.

For RNA-Seq, *A. cepistipes* was grown on MEA (for 1L MEA: 20 g malt extract, 0.5 g yeast extract, 15 g agar), RST (15 g *Picea abies* sawdust, 30 g rice, ca. 100 ml water mixed together in 720 ml jars, sterilized and overlaid with a ca. 2cm layer of homogenized tomato followed by another round of sterilization), RSTO (RST medium with additional 100 g minced orange), Orange media (3 roughly chopped oranges in 720 ml jars) and RNA was extracted using the RNEasy Midi kit (QIAGEN).

### Genome sequencing and assembly

The haploid genomes of *A. ostoyae* and *A. cepistipes* were sequenced employing the PacBio (Pacific Biosystem) RS II platform (Functional Genomics Center, ETH and University of Zurich, Switzerland; http://www.fgcz.ch). For preparing the sequencing libraries, 10 µg of gDNA aliquots were mechanically sheared to an average size of 10kb using the Covaris gTube (KBiosciences p/n 520079, Brighton, UK) in an Eppendorf microcentrifuge, and the fragment size distributions were assessed applying the Bioanalyzer 2100 12K DNA-Chip assay (Agilent p/n 5067-1508). Five μg of the sheared gDNA aliquots were DNA damage repaired, end-repaired and the final SMRTbell templates were created by blunt end ligation and exonuclease treatments. The libraries were quality inspected on an Agilent Bioanalyzer 12K DNA Chip and quantified on a Qubit.1 Fluorimeter (Life Technologies). The SMRTbell was set up by using the DNA Template Prep Kit 2.0 (3 kb - <10 kb) (Pacific Biosciences p/n 001-540-835). A ready to sequence SMRTbell-Polymerase Complex was arranged by applying the P4 DNA/Polymerase binding kit 2.0 (Pacific Biosciences p/n 100-236-500) according to the manufacturer instructions. Libraries were sequenced on 15 SMRT cell v3.0 (Pacific Biosciences p/n100-171-800), taking 1 movie of 120 minutes each per SMRT cell. The MagBead loading (PacBio p/n 100-133-600) technique served to improve the enrichment for the longer fragments. Final sequencing reports were generated for every cell, via the SMRT portal, to assess the adapter dimer contamination, the sample loading efficiency, the obtained average read length and the number of filtered sub-reads.

The HGAP3^45^ workflow of the SMRT Analysis suite v2.3 was used to create an initial assembly. After removal of redundant contigs, a scaffolding using PBJelly2^46^ and FinisherSC^47^ followed by polishing via applying the RS Resequencing protocol (SMRT Analysis suite) was performed in four iterations. The final scaffold set was checked for miss-assemblies using the RS BridgeMapper protocol (SMRT Analysis suite) and corrected if necessary. The mitochondrial scaffolds were first identified using a BLASTn search and then circularized by merging and truncating the overlapping ends. In order to correct PacBio reads, Illumina sequencing was carried out by shotgun sequencing of a 350-450 bp library with paired-end 100 bp reads to 180 fold coverage using HiSeq 2000 by the Functional Genomic Center at the ETH, Zurich. Finally, Pilon^48^ was applied to further polish the scaffolds with the genomic Illumina reads.

Draft gene models for *A. ostoyae* and *A. cepistipes* were generated by three de-novo prediction programs: 1) Fgenesh (Salamov and Solovyev 2000) with different matrices (trained on *Aspergillus nidulans*, *Neurospora crassa* and a mixed matrix based on different species); 2) GeneMark-ES (Ter-Hovhannisyan et al. 2008) and 3) Augustus (Stanke et al. 2006) with RNA-seq based transcripts as training sets. Annotation was aided by exonerate (Slater and Birney 2005) hits of protein sequences from *Armillaria* spec. to uncover gene annotation gaps and to validate de-novo predictions. Transcripts were assembled on the RNA-seq data sets using Trinity (Grabherr et al. 2011). The different gene structures and evidences (exonerate mapping, RNA-seq reads and transcripts) were visualized in GBrowse (Donlin 2009) allowing manual validation of coding sequences with a focus on Chitin, Cellulose, Pectin, Lignin, SM key genes and other genes of interest. The best fitting model per locus was selected manually and gene structures were adjusted by splitting or fusion of gene models and redefining exon-intron boundaries if necessary. tRNAs were predicted using tRNAscan-SE (Lowe and Eddy 1997). The predicted protein sets were searched for highly conserved single (low) copy genes to assess the completeness of the genomic sequences and gene predictions. Orthologous genes to all 246 single copy genes were searched for all proteomes by blastp comparisons (eVal: 10-3) against the single-copy families from all 21 species available from the FunyBASE (Aguileta et al. 2008). Additionally, the proteomes were searched for the 248 core-genes commonly present in higher eukaryotes (CEGs) by Blastp comparisons (eVal: 10-3) (Parra et al. 2009). All genomes were analyzed using the PEDANT system (Walter et al. 2009).

The genomes and transcriptomes of *A. gallica* and *A. solidipes* were sequenced using the Illumina platform at JGI. Genomic DNA was sequenced as pairs of Illumina standard and Nextera long mate-pair (LMP) libraries. For the standard libraries, 500 ng of DNA was sheared to 270 bp using the Covaris E220 (Covaris) and size selected using SPRI beads (Beckman Coulter). The fragments were treated with end-repair, A-tailing, and ligation of Illumina compatible adapters (IDT, Inc) using the KAPA-Illumina library creation kit (KAPA biosystems). For LMP, 1µg of DNA was used to generate the library using the Nextera LMP kit (Illumina). DNA was fragmented and ligated with biotinylated linkers using the Tagmentation enzyme. The fragments were circularized via ligation followed by random shearing using the Covaris LE220 (Covaris). The mate pair fragments were purified using Strepavidin beads (Invitrogen) and treated with end repair, A-tailing, and ligation of Illumina adaptors. The final product was enriched with ten cycles of PCR.

For transcriptomics, stranded cDNA libraries were generated using the Illumina TruSeq Stranded RNA LT kit. mRNA was purified from 1µg of total RNA using magnetic beads containing poly-T oligos, fragmented and reverse transcribed using random hexamers and SSII (Invitrogen), followed by second strand synthesis. The fragmented cDNA was treated with end-pair, A-tailing, adapter ligation, and ten cycles of PCR.

All libraries were quantified using KAPA Biosystem’s next-generation sequencing library qPCR kit and run on a Roche LightCycler 480 real-time PCR instrument. Except for *A. gallica* LMP, the quantified libraries were prepared for sequencing on the Illumina HiSeq sequencing platform utilizing a TruSeq paired-end cluster kit, v3(*A. gallica*) or v4(*A. ostoyae*), and Illumina’s cBot instrument to generate clustered flowcells for sequencing. Sequencing of the flowcells was performed on the Illumina HiSeq2000 or HiSeq2500 sequencer using a TruSeq SBS sequencing kit, v3 or v4, respectively, following a 2x150 indexed run recipe. Sequencing of the *A. gallica* LMP library was performed on the Illumina MiSeq sequencer using a MiSeq Reagent kit, v2, following a 2x150 indexed run recipe.

Genomic reads from each pair of libraries were QC filtered for artifact/process contamination and assembled together with AllPathsLG^49^. Illumina reads of stranded RNA-Seq data were used as input for *de novo* assembly of RNA contigs, assembled into consensus sequences using Rnnotator (v. 3.4)^50^. Both genomes were annotated using the JGI Annotation Pipeline and made available via the JGI fungal portal MycoCosm^51^.

CEGMA and BUSCO were used to assess the completeness of the assemblies. We used BUSCO Version 3.0.1 with the lineage specific profile library basidiomycota_odb9 (species:selected 41:25 with 1.335 BUSCO groups) downloaded from http://busco.ezlab.org. We performed the whole analysis in Gene set (proteins) assessment mode. Results there combined in a folder and plotted with the script generate-plot.py

### Phylogenomic analysis

We used whole-genomes of five *Armillaria* spp. and 22 Agaricales encompassing white and brown rot wood decayers, litter decomposers and ECM fungi and species from the Russulales and Boletales included as outgroups (Supplementary Table 5). An all-versus-all protein BLAST was performed using mpiBLAST-1.6.0^52^ with default parameters to find homologs based on similarity. Homologs were clustered into protein families using the Markov Cluster Algorithm^53,54^ with an inflation parameter of 2.0. For each cluster, a multiple sequence alignment was inferred using PRANK v.150803 (Loytynoja A et al, 2005), run with default settings. Subsequently, spuriously aligned regions were excluded with trimAl v1.4.r15 (Capella-Gutierrez et al, 2009), with a gap threshold of 0.2. Next, we inferred Maximum likelihood (ML) gene trees for each cluster of a minimum of four proteins using FastTree v2.19 (Price et al, 2010). FastTree was run with CAT/GAMMA20 model for rate heterogeneity and an improved LG model^55^ for substitution probabilities. Gene trees were mid-point re-rooted using Phyutility v 2.2.6^56^.

Next, we screened the gene trees with a custom Perl script^57^ for the presence of deep paralogs and inparalogs. Gene trees with deep paralogs were eliminated, while in trees with inparalogs, the paralog closest to the root was retained in the corresponding alignments. Finally, 835 clusters passed these criteria and had >50 amino acid sites and (>=) 60% taxon occupancy were concatenated into a supermatrix with 188895 sites. We used this supermatrix to infer species tree with RAxML PTHREADS version 7.2.8^58^ using a partitioned WAG+G model, where each data partition represented a single input gene family. To assess branch support of the tree we performed a bootstrap analysis in RAxML with 100 replicates under the same model.

We used penalized likelihood (PL) implemented in the program r8s version 1.70^59^ for molecular clock dating based on the optimal ML tree and two fossils and one secondary calibration. *Archaeomarasmius legettii* (94-90 Myr)^60^ was used to define minimum age constraint (92 Myr) for the origin of marasmioid clade (*sensu* Matheny 2006^61^). *Palaeoagaricites antiquus* (100-110 Myr) is the oldest fossil can be placed within Agaricales^62^, hence we constrained the most recent common ancestors (mrca) of this order to 108 Myr minimum age. To define the minimum age constraint of the origin of Boletales to 84 Myr, we referred to the analyses of Kohler *et al*.^27^. Maximum age constraints were defined with a wide safety window to allow for the calibrated clades to be inferred at least twice as old as their minimum ages. The mrca of the Boletales, that of the Agaricales and that of the marasmioid clade were constrained between 84-250 Myr, 108-250 Myr and 92-180 Myr, respectively, as minimum and maximum ages. A cross-validation (CV) analysis was used to identify an optimal smoothing parameter between 10^-20^ and 10^9^. All analyses were run in four replicates (random number seeds). We applied additive penalty function and run optimization for 25 times initialized from independent starting points. Furthermore after reaching an initial solution of an optimization step, the solution was perturbed and the truncated Newton (TN) optimization was rerun 20 times. We found that the optimal smoothing parameter (λ) can be between 10^-7^ and 10^-3^, thus we estimated divergence time with five lambda parameter ranges between 10^-7^ and 10^-3^. By checking the gradients and the estimated ages we found a high similarity between the results of the analyses obtained with different smoothing parameters (Supplementary Fig. 2). By inspecting the CV analyses and the gradient values a smoothing parameter of 10^-6^ was chosen as the most adequate and was used to estimate divergence times.

### COMPARE analysis

We analyzed the evolutionary history of gene families of *Armillaria* species and closely related Agaricales using the COMPARE pipeline^63^. To reconstruct gene duplications and losses, a genome-wide collection of 13821 gene trees (see previous section) was first reconciled with the species tree using Treefix v1.1.10^64^. Treefix was run with RAxML site-wise likelihood model, Maximum Parsimony Reconciliation model (MPR) and an --alpha threshold of 0.001 to find any alternate gene tree topologies that minimize duplication/loss cost but are statistically equivalent to ML gene tree. Orthologs were identified and recoded into presence/absence matrix, as described in Nagy et al.^63^. We then inferred duplications and losses for each orthogroup along the species tree using Dollo parsimony. Gene trees with less than four proteins were excluded. Subsequently, we also reconstructed the genome size for a given node by summing over gains and losses to the genome size of most recent common ancestor (MRCA). GO enrichment analysis based on the Fisher exact test with Benjamin-Hochberg correction was performed using Pfam domains mapped to protein clusters and creating GO annotations with pfam2go version 2016/10/01^65^ followed by enrichment analysis at p<0.05.

### Copy number analyses

We analyzed the copy number distribution of selected gene families in *Armillaria*, *Moniliophthora*, and *Heterobasidion annosum* and, non-pathogenic members of Agaricales. For this, predicted proteomes of all the taxa under study were annotated with Pfam domains and IPR signatures. We scanned the protein sequences for pfams using pfamscan.pl v1.5 against the Pfam 30.0 database and InterPro signatures using InterProscan v5. Next, we built search terms for 42 PCWDE families and 25 putative pathogenicity related genes based on literature evidence (‘Search_Terms’ in Supplementary Table 5). In few cases, Pfam signatures were either absent or did not yield consistent results. CBM67 copy numbers were obtained based on BLAST-hits by counting homologs obtained through blasting proteins annotated by JGI as CBM67. Genomes were searched iteratively for significant hits to avoid the impact of phylogenetic distance on the detection of homologs. We annotated cellobiose dehydrogenases (CDH, AA3_1), alcohol oxidases (Aox, AA3_3) and GMC oxidoreductases (GMT, AA3_2) using a tree-based approach. To this end, proteins of *Armillaria gallica* (Armga1) were combined with homologs in other genomes, followed by multiple sequence alignment construction using MAFFT v7.273^66^ with -auto option and the estimation of a gene tree with FastTree v2.19 (as above). Classification was done based on the occurrence as monophyletic groups in the phylogenetic tree. Small secreted proteins (SSPs) were defined as proteins with less than 300 amino acids and sequence-based evidence for secretion as inferred by signalP v4.1^67^. Both signalP-noTM and signalP-TM models were used for the detection of signal peptides.

### Analysis of transposable elements

#### De novo element discovery and annotation –

We used the REPET package version 2.5^68,69^ to identify, classify and annotate transposable elements (TEs) within the genomes studied. Since the *A. mellea* genome contained large amounts of “chaff” contigs, we thresholded this assembly to only include contigs larger than 1000 bps prior to further analysis.

The REPET *de novo* pipeline was run using genome self-alignment as well as a search for structural features using LTRHarvest^70^. Consensus sequences were clustered using Piler v1.0^71^, Recon 1.08^72^ and Grouper^73^ and classified using PASTEC^69^. The resulting consensus sequences were filtered for low frequency elements, short sequence repeats and sequences that were classified as putative host genes and then clustered into families using Markov clustering^53^. For annotation, the consensus sequences collected from each species were combined into a pan-species TE library. TE identification was then carried out using the REPET anno pipeline, implementing BLASTER^73^, RepeatMasker 4.06 and CENSOR 4.2.29^74^. Configuration files for the pipelines containing detailed parameter settings as well as a sample command line script can be found at https://github.com/JackyHess/Armillaria_TE_annotations

#### TE organization along the genome in core *Armillaria* species –

In order to gain an understanding of how TEs are organized among the core *Armillaria* genomes we investigated TE and gene content density in 50 kb genome windows. Each genome was segmented into 50 kb partitions using bedtools (http://bedtools.readthedocs.io) and for each partition, TE coverage and genic coverage were estimated. Prior to estimating genic coverage, we filtered annotated CDS for which 20% of the sequence overlapped with TE annotations to remove putative TE-derived genes. This reduced the number of protein-coding genes considered by up to 7000 (Supplementary Table 1). All analysis scripts can be found at https://github.com/JackyHess/Armillaria_TE_annotations.

### In-depth transcriptome analysis of A. ostoyae

Whole transcriptome sequencing was performed using TrueSeq RNA Library Preparation Kit v2 (Illumina) according to the manufacturer’s instructions. Briefly, RNA quality and quantity measurements were performed using RNA ScreenTape and Reagents on TapeStation (all from Agilent) and Qubit (ThermoFisher); only high quality (RIN >8.0) total RNA samples were processed. Next, RNA was DNaseI (ThermoFisher) treated and the mRNA was purified and fragmented. First strand cDNA synthesis was performed using SuperScript II (ThermoFisher) followed by second strand cDNA synthesis, end repair, 3’-end adenylation, adapter ligation and PCR amplification. All purification steps were performed using AmPureXP Beads (Backman Coulter). Final libraries were quality checked using D1000 ScreenTape and Reagents on TapeStation (all from Agilent). Concentration of each library was determined using the QPCR Quantification Kit for Illumina (Agilent). Sequencing was performed on Illumina NextSeq instrument using the NextSeq Series 2x150bp high-output kit (Illumina) generating >20 million clusters for each sample.

Bioinformatic Analysis Draft genome sequence together with genome annotation file was used as a reference for *A. ostoyae* RNA-Seq analysis. Paired-end Illumina NextSeq reads were quality trimmed using CLC Genomics Workbench tool version 9.5.2 (CLC Bio/Qiagen) removing ambiguous nucleotides as well as any low quality read end using an error probability cutoff value of 0.05 (corresponding to Phred score 13). Trimmed reads were mapped using the RNA-Seq Analysis 2.1 package in CLC allowing intergenic read mapping and requiring at least 80% sequence identity over at least 80% of the read lengths; strand specificity was omitted. Reads with less than 30 equally scoring mapping positions were randomly mapped; reads with more than 30 potential mapping positions were considered as uninformative repeat reads and were excluded from the analysis (Supplementary Table 6).

“Total gene read” RNA-Seq count data was imported from CLC into R version 3.0.2. Genes were filtered based on their expression levels keeping only those features that were detected by at least five mapped reads in at least 25% of the samples included in the study. Subsequently, “calcNormFactors” from package “edgeR” version 3.4.2^75^ was used to perform data normalization based on the “trimmed mean of M-values” (TMM) method^76^. Log transformation was carried out by the “voom” function of the “limma” package version 3.18.13^77^. Linear modeling, empirical Bayes moderation as well as the calculation of differentially expressed genes were carried out using “limma”. Genes showing at least four-fold gene expression change with an FDR value below 0.05 were considered as significant. Multi-dimensional scaling (“plotMDS” function in edgeR) was also applied to visually summarize gene expression profiles revealing similarities between samples. In addition, unsupervised cluster analysis with Euclidean distance calculation and complete-linkage clustering was carried out on the normalized data using “heatmap.2” function from R package “gplots”.

### Proteomics

#### Whole cell lysate preparation and trypsin digestion

Two hundred mg (±5%) from every developmental stage of *A. ostoyae* was ground in 600 µL of ice-cold lysis buffer (100 mM Tris-HCl, 50 mM NaCl, 20 mM EDTA, 10 %(v/v) glycerol, 1 mM PMSF and 1 µg/mL Pepstatin A pH 7.5) using a IKA T10 basic homogenizer (IKA-Werk GmbH & Co. KG, Germany) and PT 1200 E Ergonomic Homogenizer (POLYTRON PT 1200 E; Kinematica AG, Switzerland), respectively. Samples were sonicated (Bandelin Sonoplus HD2200, Bandelin Elec., Germany) three times on ice (MS73 probe, Cycle 6, 10s, Power 20%), lysates were incubated overnight on ice at 4°C and then clarified by centrifugation at 9,700 g for 10 min. Protein was quantified using the Bradford assay and samples were normalized for protein concentration, where possible, per replicate. Samples were brought to 15% (w/v) trichloroacetic acid (TCA) for precipitation, with acetone washes. Samples were resuspended in 6M urea, 2M thiourea and 100mM Tris-HCl pH 8.0 (60-120 µL), protein concentrations analyzed, as above, and adjusted to 1 M urea final using ammonium bicarbonate prior to trypsin digestion. Protein samples (10 µg each) were sequentially reduced and alkylated with 5 µM DTT and 15µM IAA, respectively^11^, brought to 0.01% (v/v) ProteaseMAX and trypsin added (1.6µL; 1 µg/µL) to a final protein amount (10µg) (37°C, overnight). Samples were acidified by addition of 1µL trifluoroacetic acid (TFA). Peptide solutions were then dried in a centrifugal evaporator, resuspended in 0.5% (w/v) TFA and desalted using C_18_ Ziptips (Millipore^â^ Ziptips C18)^78–81^.

#### LC-MS, data processing and interpretation

Peptide mixtures were analyzed using a Thermo Fisher Q-Exactive^TM^ mass spectrometer coupled to a Dionex RSLCnano for LC-MS/MS analysis. LC gradients operated from 3-40%B over 40 min, with data collection using a Top15 method for MS/MS scans^82^. LC-MS spectra chromatograms were analysed manually using rawMeat software. Raw files corresponding to the aforementioned spectra were then processed against an *A. ostoyae* predicted protein database using MaxQuant (version 1.3.0.5) and further filtered and visualized in Perseus (Version1.4.1.3)^82^. MaxQuant parameters were as described in Owens *et al.^81^*. Peptide intensity values were normalized to log2 values in Perseus. Only samples represented in all replicates of a sample group were taken (n = 1 to 4). Proteins common to all stages were z –score normalised, averaged and hierarchical clustering was performed to generate heat maps and profile plots. Venn diagrams were prepared for all proteins (unique and abundant proteins inclusive) in the entire data set (http://bioinformatics.psb.ugent.be).

Proteomic analysis revealed the presence of 2,549 proteins, whereby 39.7% (1012/2,549) proteins were common to all *A. ostoyae* developmental stages and 25.2% (643/2,549) were unique to individual developmental stages. Proteins were unique to rhizomorph stage (RH; *n* = 286), vegetative mycelia (VM; *n* = 163), young fruiting body (YFB: *n* = 97), primordial stages 1 and 2 (PS1,2; *n* = 73) and mature fruiting body (MFB; *n* = 24). Quantitative changes in protein abundance were also evident between different development stages, albeit to different extents. For instance, comparing RH to VM stages, individual proteins underwent increased (*n* = 184) and decreased (*n* = 125) abundance (total detected *n* = 2190), whereas comparative analysis of YFB against MFB revealed only eight proteins with increased abundance (total detected *n* = 1203). Comparative analysis of all other developmental stages yielded intermediate changes in the abundance of specific proteins.

### Motif Discovery

We used 1 kb sequences of co-expressed genes (which most likely contain their promoter), located upstream of the start codon, including 5’ UTR, to predict putative TFBS. We performed *de novo* motif discovery in our co-expressed gene sets using Weeder 2.0^39,83^. We created frequency files from promoter regions of all genes of *A. ostoyae* using the Weeder2.0 frequencymaker. Frequencies were counted on both strands for motif lengths 6, 8 and 10, allowing 1, 2 and 3 mismatches, respectively. Using the inferred frequency counts as reference, both strands of the sequences were scanned using Weeder, allowing the detection of a maximum of 100 motifs. Next, we grouped the motifs based on Pearson correlation co-efficient, aligned using the ungapped Smith-Waterman algorithm and clustered using UPGMA, all executed via the STAMP webtool^84^. Finally, we constructed familial binding profiles for each motif group and searched for matching DNA motifs in the JASPAR Core (Fungi) 2016 database using the TOMTOM webserver^85^.

### Data availability

Genome assemblies and annotation were also deposited at DDBJ/EMBL/GenBank under the following accessions *A. ostoyae*: FUEG01000001-FUEG01000106, *A. cepistipes*: FTRY01000001-FTRY01000182, *A. gallica*: NKEW00000000and *A. solidipes*: NKHM00000000. Gene Expression Omnibus (GEO) archive of the sequenced A. ostoyae libraries was deposited in NCBI’s GEO Archive at http://www.ncbi.nlm.nih.gov/geo under accession GSE100213.

